# Weak form Scientific Machine Learning for Systems Biology: A Tutorial on WENDy

**DOI:** 10.64898/2026.07.02.735880

**Authors:** Nora Heitzman-Breen, Rainey Lyons, Paras Jain, Mohit K. Jolly, David M. Bortz

## Abstract

Mechanistic ordinary differential equation models are widely used in systems biology to represent biochemical networks, population dynamics, cell-state transitions, and other biological processes; however, their predictive value depends critically on accurate parameter estimation from noisy and often sparse experimental data. In this tutorial, we present the Weak-form Estimation of Nonlinear Dynamics (WENDy) method as a forward-solver-free approach that reformulates parameter estimation as a covariance-corrected weak-form regression problem by integrating the model equations against compactly supported test functions. We present the background on the methodology through the lens of the familiar logistic equation, and we demonstrate applications of the method on real experimental data through two systems biology examples: a glycolytic oscillator with relatively dense time-course data and a sparse epithelial-mesenchymal cellstate transition model with multiple experimental replicates. Ultimately, using WENDy, we estimate interpretable biological parameters with uncertainty for systems with noisy and sometimes sparse available experimental data.

## 1 Introduction

A central goal of systems biology is to understand how biological function emerges from networks of interacting molecular, cellular, and population-level processes [1]. Mathematical models provide a principled way to encode such interactions as quantitative hypotheses. In this setting, parameter estimation is not simply a numerical post-processing step, but rather a step that connects proposed mechanisms to experimental data. This connection is what determines whether the hypothesized model can make reliable and biologically interpretable predictions.

Many models in systems biology can be formulated as systems of ordinary differential equations (ODEs),

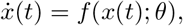

where *x*(*t*)*∈* ℝ^*d*^ denotes a vector of measured or latent states and *θ* denotes a vector of parameters. Such models have been used to study glycolysis [2–5], cell signaling [6–9], circadian rhythm [10, 11], and cell population dynamics [12, 13]. Given experimental observations, the goal of parameter estimation methods is to find an appropriate *θ* such that the model trajectory agrees (in some sense) with recorded data. In practice, this is most often posed as an output-error, maximum likelihood, or Bayesian inverse problem in which the ODE model is repeatedly solved for many trial parameter values [14, 15]. This forward-solver based philosophy is a natural and well-studied approach which has shaped much of the computational infrastructure of systems biology.

However, this forward-solver structure also introduces significant computational and statistical challenges. Each single objective function evaluation requires solving the ODE system. Moreover, gradient-based methods often require adjoint or sensitivity calculations. For stiff, multiscale, or high-dimensional systems, these hurdles can accumulate into a high computational cost. Additionally, errors associated with experimental data can compound these challenges. Biological data, in particular, can be noisy, partially observed, or sampled irregularly or sparsely.

There have been several alternatives to repeated forward simulations proposed in the literature. Generalized profiling methods replace direct trajectory matching with a smoothed representation of the state trajectory and enforce the ODE system as a penalty or residual constraint [16]. Related gradient-matching and least-squares-type methods estimate derivatives from smoothed data and then fit the right-hand side of the ODE [17]. These methods can offer substantial speedups, but they introduce choices of smoothing parameters, derivative approximations, and regularization weights that can strongly influence performance in the presence of noise.

Weak-form Scientific Machine Learning (WSciML) provides a complementary route. Instead of estimating derivatives from the data or repeatedly solving the ODE system, weak-form methods utilize the eponymous weak-form to transfer derivative calculations from noisy data onto analytically known test functions, producing algebraic equations involving only integrals of the observed state and nonlinear model features. These test functions also play a role similar to a Gaussian smoother and can enable accurate parameter recovery even at high noise levels [18]. Leveraging test functions, also sometimes called modulating functions, in system identification was first proposed by Shinbrot [19] and further explored by Loeb and Cahen [20]. Two key methods in WSciML recently developed within our group include the Weak-form Sparse Identification of Nonlinear Dynamics (WSINDy) method for model discovery [21, 22] and the Weak-form Estimation of Nonlinear Dynamics (WENDy) [23] method for parameter estimation, the focus of this tutorial. When compared to classical forward-solver approaches, WENDy has been shown to consistently be faster and more accurate at parameter estimation for many systems [23–25]. For an overview of this novel class of methods, we direct the interested reader to [26, 27].

The purpose of this tutorial is to make WENDy accessible to systems biology researchers who routinely fit mechanistic ODE models to experimental data. We present the weak-form construction, explain the role of test functions and noise models, and demonstrate how the method is implemented in code and how uncertainty estimates are generated. Our goal is not to argue that weak-form methods completely replace forward simulation-based methods. Forward solves remain essential for model validation, prediction, experimental design, and comparison of fitted trajectories to observed data. Rather, WENDy provides an efficient parameter estimation mechanism that can reduce the computational burden of calibration and make it easier to explore candidate models, initial guesses, constraints, and uncertainty estimates. For systems biology applications where data are noisy, sparse, and expensive to collect, this weak-form perspective offers a useful addition to the existing parameter estimation toolkit.

The remainder of this manuscript is structured in the following way. In Section 2, we demonstrate the method on two systems biology examples. The first example considers a glycolytic oscillator, representing a setting with relatively dense time-course data and nonlinear oscillatory dynamics. The second example considers a sparse cell-state transition model for epithelial–mesenchymal heterogeneity, in which only a small number of experimental time points are available across multiple initial culture conditions. In Section 3, we provide a discussion of the results and of some caveats of the method. In Section 4, we provide the details of the WENDy algorithm from the lens of a simple logistic model. We also include artificial experiments on this model showcasing the WENDy method’s performance as the noise level increases and the data sampling coarsens. For a general description of the method, we will direct readers to the works [23–25].

## 2 Results

In this section, we apply WENDy to perform parameter estimation on two systems biology examples and provide a tutorial on the implementation of the WENDy algorithm in MATLAB. Firstly, we demonstrate WENDy in application to single-experiment data with a high collection frequency and minimal measurement noise. Then, we demonstrate WENDy in application to multi-experiment data with a low collection frequency and measurement noise.

### 2.1 Example 1: Parameter Estimation of Glycolytic Oscillator

Temporal oscillation of key metabolites of glycolysis can be observed in yeast cells [28–32]. Two such metabolites are NADH and ATP. For this example, we will model these two glycolytic metabolites using the two-compartment cubic order equations given below,

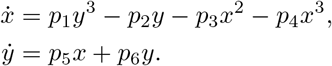

Here, *x* represents fluorescence of NADH and *y* represents fluorescence of ATP. This model is inspired by the cubic model in the original work [2] which was used to successfully model NADH and ATP reaction dynamics.

#### Data Entry

For this example, we use shifted normalized NADH and ATP fluorescence data generated in [33] and publicly available in [34]. NADH was measured via fluorescence, and ATP was measured via amper-based nanosensor [33]. For this demonstration, we select the first three periods present in the data to perform parameter estimation on model (2.1). The data consists of N=10,000 observations from a single realization of an experiment, where each observation is comprised of two normalized flourescence intensities corresponding to the model compartments for NADH and ATP. We create a `wsindy_data` object from the experimental observations as shown in Listing 1. Here, by default, the `wsindy_data` object will assign an additive Gaussian error structure to the data, which informs the covariance correction as discussed in Section 4.

**Listing 1.**
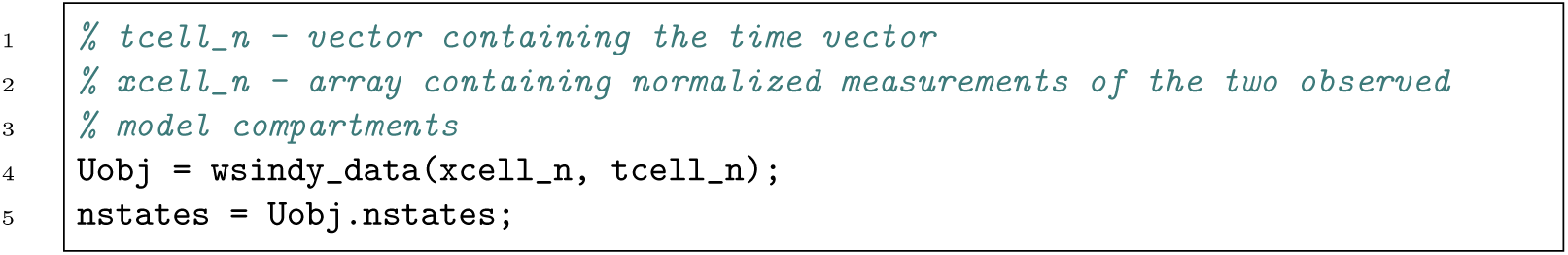
Creating a single `wsindy_data` object.

#### Equation Creation and Regression

Next, we construct a library of model features. We have two options for constructing this library: either calling predefined library terms or manually entering the library terms. For this example, we will call predefined library terms to reconstruct model (2.1). For guidance on manual entry of model terms, see Section 2.2. We assume that all model parameters *p* ={*p*_1_, *p*_2_, *p*_3_, *p*_4_, *p*_5_, *p*_6_} are unknown. Each term in model (2.1) that is associated with a parameter in *p* will be a feature in the library, and because each term is polynomial, we can call a predefined library term using a vector of the polynomial degree of each state variable. For example, the predefined library term, *x*^4^*y*^2^*z*^9^, is called by the vector [4, 2, 9], which contains the polynomial order of the state variables *x, y*, and *z*, respectively. We create the feature library corresponding to model (2.1) in Listing 2. Note that the left-hand side of the model equations is assumed to be first-order derivatives of state variables unless otherwise specified. See Section 2.2 for creation of an alternative left-hand side equation.

**Listing 2.**
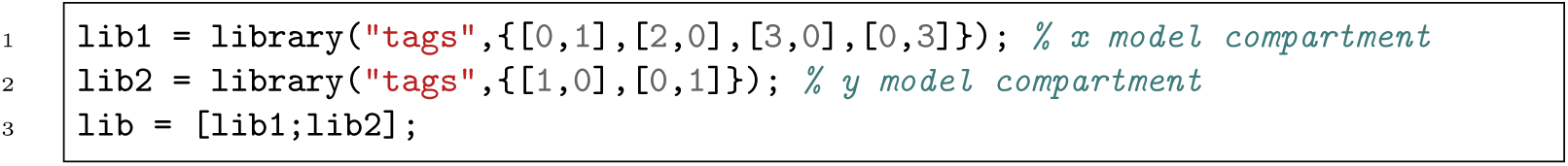
Creating right-hand side library object.

Lastly, the `wsindy_data` and library objects are combined in a regression model `wendy model` and solved using the WENDy algorithm. The `wendy_model` builds the weak-form linear system (7) as described in Section 4.

For convenience, we provide a `WENDy_defaults.m` script which handles any hyperparameters and the test function object creation. We provide more detail on test function customization in Section A. The `wendy` solver then performs the (IRLS) algorithm to produce the coefficient vector `WS.weights` = [*p*_1_, *p*_2_, *p*_3_, *p*_4_, *p*_5_, *p*_6_]. The code for this is provided in Listing 3. The function `WS_opt.wendy()` performs the WENDy method on the `wendy_model` object. While we make use of only the edited model object and the parameter covariance matrix `CovW`, the function also outputs other variables which can be used for troubleshooting. We describe these outputs in Table B2.

**Listing 3.**
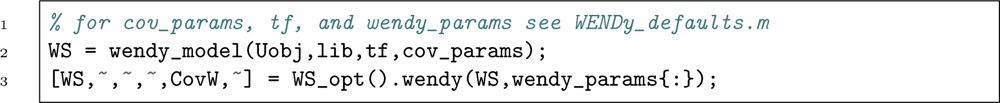
Creating `wendy_model` and applying the WENDy method.

#### WENDy Outputs and Confidence Interval Generation

The WENDy algorithm provides parameter estimates along with the parameter covariance structure, from which confidence intervals can be easily calculated. Figure 1 shows the dynamics of model (2.1) with parameters found using the WENDy algorithm and normalized NADH-ATP fluorescence data [33, 34] match the experiment well.

**Fig. 1.**
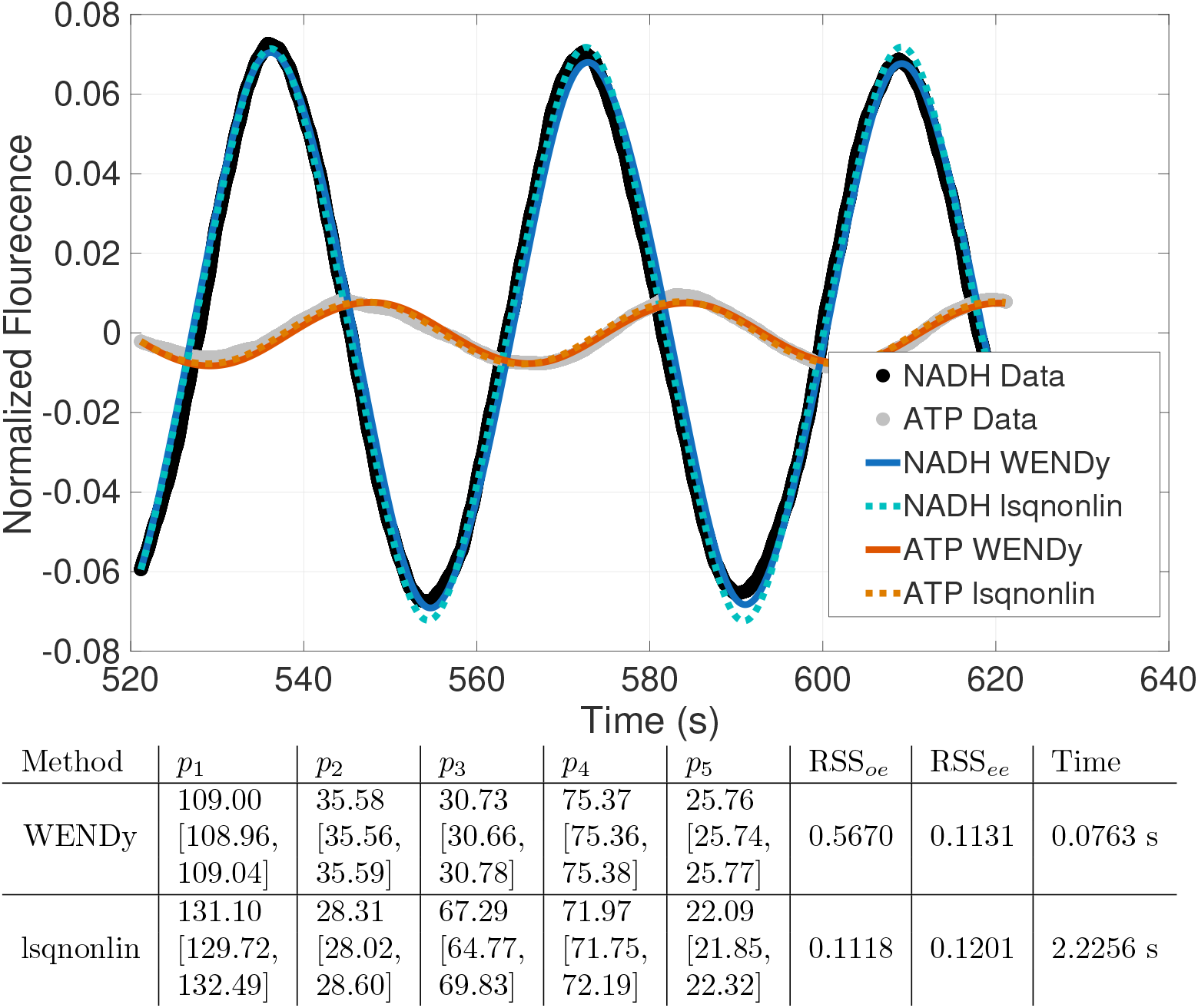
Model (2.1) dynamics for parameters *p* = {*p*_1_, *p*_2_, *p*_3_, *p*_4_, *p*_5_, *p*_6_} found using WENDy describes glycolitic oscillations well and performs faster compared to the output error based method. (Top) Normalized NADH and ATP fluorescence over 120 minutes from data generated in [33]. Data was obtained from the [34]. Dynamics given by the model (2.1) for NADH (blue lines) and ATP (red lines) using parameters found from WENDy (solid lines) and from lsqnonlin (dotted lines) versus normalized experimental fluorescence data for NADH (black dots) and ATP (grey dots). (Bottom) Table of parameter estimates and corresponding 95% confidence intervals.

For comparison, we also consider the model fit using the `lsqnonlin` function in MATLAB [35] with default settings. Both the WENDy and `lsqnonlin` parameter estimates along with their respective 95% confidence intervals are listed in Figure 1. We additionally calculate the residual sum of squares considering both the output error (RSS_*oe*_) and (weak-form) equation error (RSS_*ee*_) residuals. Note that, as expected, the RSS_*ee*_ is lower for the WENDy parameter estimate, as this method is based on optimization of equation error. The RSS_*oe*_ is lower for the `lsqnonlin` parameter estimate, as this method is based on optimization of output error. In general, the parameter estimates are similar between the two methods and produce similar model dynamics, as demonstrated in Figure 1. However, for this oscillatory system, the WENDy algorithm is *∼*3 times faster than the default `lsqnonlin` implementation, 2.43 s versus 7.46 s. It should also be noted that the `lsqnonlin` method was initialized with the recovered WENDy parameters to avoid convergence failures of the method.

### 2.2 Example 2: Parameter Estimation of Cell State Transition Model

For this example, we consider two population models of epithelial-mesenchymal cancer cell state transitions. We first model the ephithelial cell fraction (*E*) and mesenchymal (*M*) fraction using the growth and transition (G&T) model from [12] is given below,

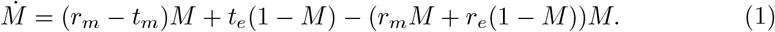

Here, *r*_*m*_ represents the mesenchymal cell growth rate per unit time, *t*_*m*_ represents the mesenchymal-to-epithelial transition rate per unit time, *t*_*e*_ represents the epithelial-to-mesenchymal transition rate per unit time, and *r*_*e*_ represents the epithelial cell growth rate per unit time.

We additionally consider the growth, influence, and transition (GI&T) model from [12], which is given below,

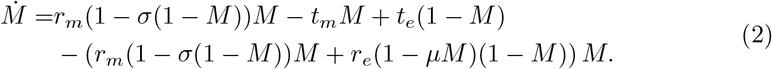

Here, *σ* and *µ* are unitless parameters that represent the strength of reduction in mesenchymal or epithelial cell growth in the presence of the other cell population. Note that we assume all cells are either epithelial or mesenchymal, i.e. 1 = *E* + *M*.

#### Data Entry and Interpolation

The data used in this example contains three observations of two experiments, giving a multi-trajectory WENDy problem with six trajectories (three realizations *×* two initial culture conditions). The data is observed as cell fractions, where Culture 1 starts with a high initial mesenchymal fraction (assumed *M*_0_ = 0.999), and Culture 2 starts with a low mesenchymal fraction (*M*_0_ = 0.001). Each realization of the experiment occurs two weeks apart over an 8-week time frame. To apply the WENDy method in this low collection frequency system, we interpolate the sparse data to provide the necessary state data required to form the weak-form integrals. More information on the validity of pre-processing sparse data through interpolation in conjunction with the WENDy algorithm is provided in Section 4. We call the realizations of the experiment trajectories, each of which becomes its own `wsindy_data` object, and are then combined into an array (see Listing 4). This array of data objects prompts WENDy to consider multiple trajectories. In this data object, we pass the estimated noise type (multiplicative lognormal in this case), which informs the covariance correction discussed in Section 4.

#### Equation Creation and Regression

Once the data object array is created, we then construct the library features. In [12], there are multiple models considered, but for the sake of the tutorial, we consider the

**Listing 4.**
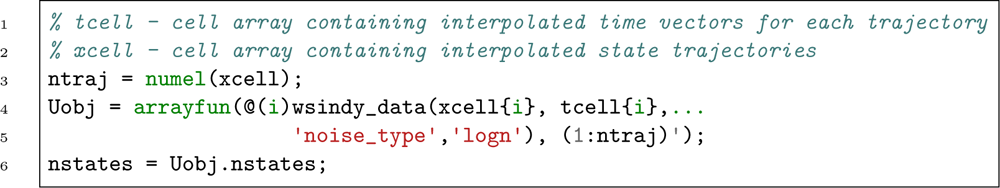
Creating `wsindy_data` object array.

G&T model for mesenchymal fraction:

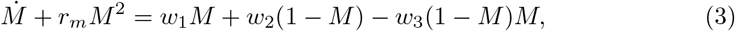

where *r*_*m*_ is a known proliferation rate fixed using the same value as in [12] and **w** := (*w*_1_ = *t*_*m*_, *w*_2_ = *t*_*e*_, *w*_3_ = *r*_*e*_)^*T*^ are the unknown parameters to be learned. Note that, unlike in Section 2.1, we cannot use the default left-hand side for this more complex model as the known terms (*r*_*m*_*M* ^2^ in this case) are moved to the left-hand side. In Listing 5, we provide the code to specify the left-hand side object. We once again utilize the predefined library terms. For example, the term 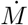 is a first-order polynomial and the only state variable, and is therefore called as a function by using the tag [1]. Here, `LHSterm1` has an additional property linOp = [1], indicating the first derivative. Lastly, `LHSterm2` has a coefficient property `coeff` since *r*_*m*_ is a known value in this example.

**Listing 5.**
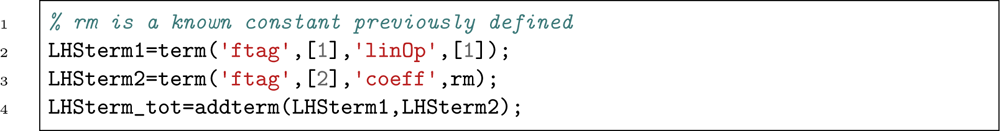
Creating left-hand side library object.

For this example, we will construct the library of features by manually defining terms rather than using predefined terms. We will need to create a corresponding library term for each term in the right-hand side of model (3) (i.e. *M*, (1*− M*), and (1 *− M*)*M*). These definitions are made in Listing 6, and are then combined into the feature library object.

As done in Section 2.1, the objects are used to construct the regression model `wendy model`, which is then solved using the WENDy algorithm. During the application of the WENDy algorithm, the data is concatenated across trajectories, stacking all subsystems into a single overdetermined system. Additionally, for this example, we wish to consider biologically informed bounds on the parameter ranges considered for the estimation algorithm. Upper and lower bounds can be provided as vectors with the same size as the unknown parameters. Listing 7 calls the WENDy algorithm with parameter constraints.

**Listing 6.**
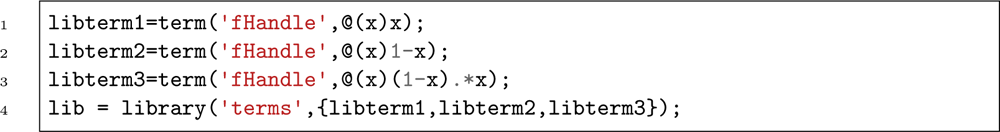
Creating right-hand side library object.

**Listing 7.**
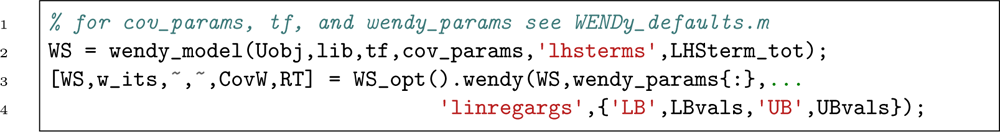
Creating `wendy_model` and applying the WENDy method.

#### WENDy Outputs and Confidence Interval Generation

The WENDy algorithm provides parameter estimates. Figure 2 shows that the dynamics of models (1) and (2) with parameters found using the WENDy algorithm and mesenchymal cell fraction data match the experiment well.

**Fig. 2.**
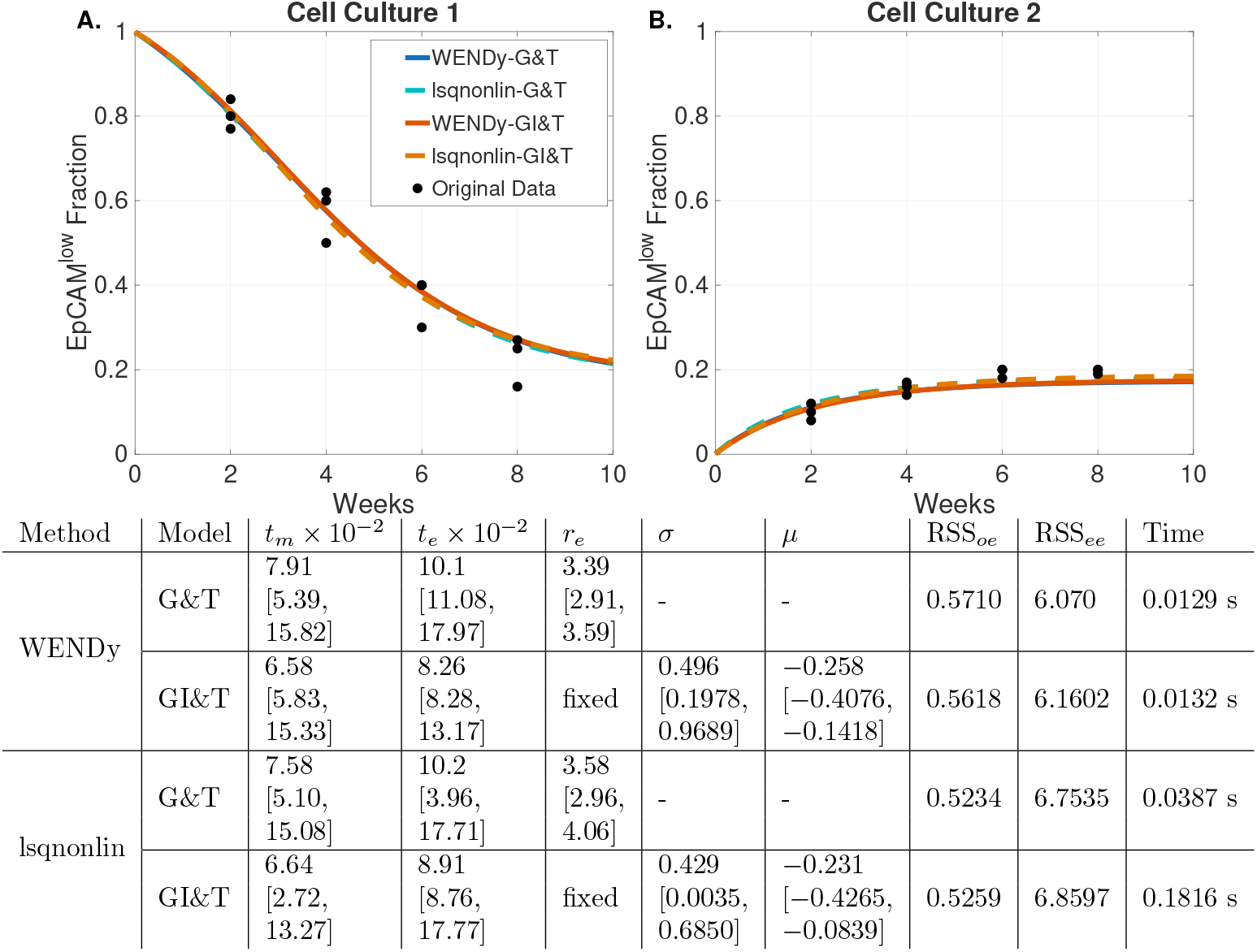
G&T and GI&T models dynamics for parameters found using WENDy describe mesenchymal cell populations well and are consistent with an output error based method. (Top) Mesenchymal cell populations over 8 weeks for experiments where the initial E-M ratio is (A.) low or predominantly mesenchymal and (B.) high or predominantly epithelial. Dynamics given by the G&T model (1) using parameters found from WENDy (solid blue line) and from lsqnonlin (dotted blue line) and given by the GI&T model (2) using parameters found from WENDy (dashed red line) and from lsqnonlin (dashed red line) versus experimental data (black dots) from [36]. (Bottom) Parameter estimates and corresponding 95% confidence intervals. For both models, we fix *r*_*m*_ = 3.1 per week, and in the GI&T model, we fix *r*_*e*_ = 4.8 per week.

For comparison, we also consider the model fit using the `lsqnonlin` function in MATLAB [35] with default settings. Both the WENDy and `lsqnonlin` parameter estimates, along with their respective 95% confidence intervals, are listed in Figure 2. For this example, we calculate confidence intervals by parameteric bootstrapping to avoid overstating certainty in the parameter estimates driven by the addition of data points through the interpolation process. As before, we additionally calculate the residual sum of squares considering both the output error (RSS_*oe*_) and weak-from equation error (RSS_*ee*_) residuals. Once again, the RSS_*ee*_ is lower for the WENDy parameter estimate, and the RSS_*oe*_ is lower for the `lsqnonlin` parameter estimate. In general, the parameter estimates are similar between the two methods and produce similar model dynamics, as demonstrated in Figure 1. Furthermore, both the output error and equation error residuals are in agreement that the GI&T model fits the data better than the G&T model, as found in [12]. Here, the WENDy algorithm performs similarly to the default `lsqnonlin` implementation in terms of time, 0.5 s versus 1 s.

## 3 Discussion

In this tutorial, we presented the WENDy method as a forward-solver-free approach for parameter estimation for models arising in systems biology. The examples considered illustrate two distinct regimes in which weak-form parameter estimation can be useful. In the glycolytic oscillator example, the data is relatively dense in time, and the underlying oscillatory dynamics are nonlinear and numerically stiff. In this setting, WENDy provides an efficient alternative to output-error methods as the stiff nature of the model causes instabilities in the parameter estimates and can even cause `lsqnonlin` to fail to converge. It is worth noting that this failure of forward solver methods can be partially mitigated and convergence can be substantially accelerated by using the recovered WENDy parameters as the initial guess in forward-solver methods, demonstrated previously in [23, 24]. On the other hand, in the cell state transition example, we demonstrate that WENDy can be used in a low-data setting when the dynamics are sufficiently smooth, and interpolation is a reliable method for filling out the weak-form integrals.

A common theme across both examples is that WENDy works well for models who are *linear in parameters*. This includes many polynomial, rational (after rearrangements), mass-action, and compartmental transition models. Generally, WENDy works well for any model that can be written in the form

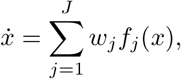

where the terms *f*_*j*_(*x*) can be nonlinear. What is important for the standard WENDy formulation is not that the model is linear in the state *x*, but linear in the unknown parameters *w*_*j*_. This makes WENDy particularly well matched to many interpretable systems biology models where biological mechanisms are encoded through prescribed terms and the goal is to estimate rates, interaction strengths, or transition parameters.

The sparse cell-state transition example also highlights an important practical point. Weak-form methods avoid numerical differentiation of noisy data, but they do not eliminate the need for adequate temporal information. When the data is sparse, the weak-form integrals must still be approximated with low quadrature error from the available measurements. If the dynamics are smooth and slowly varying, as in the cell-state transition model, linear interpolation provides a reasonable reconstruction for quadrature (more detail is provided in Section 4.2). However, interpolation should not be interpreted as generating new independent data. Interpolated points are deterministic functions of the original observations and inherit strong correlations. Consequently, uncertainty estimates based on treating interpolated points as independent measurements can be misleading. For this reason, the confidence intervals are obtained by bootstrapping the original sparse observations and repeating the interpolation-WENDy estimation pipeline.

Despite these advantages, several WENDy-specific limitations should be kept in mind. The standard formulation assumes that the governing equations can be expressed as a linear combination of known feature functions with unknown coefficients; consequently, models involving unknown Hill coefficients, unknown delay times, nonlinear parameter dependencies, or genuinely nonlocal terms require novel extensions of WENDy. In addition, the version of WENDy presented in this work assumes that all dynamically relevant state variables are observed, or at least can be reconstructed with sufficient accuracy. If important compartments or latent variables are unobserved, then direct weak-form regression on the measured variables alone can produce biased estimates because the fitted working model is incomplete. Addressing these issues is an active research direction, with foundational progress reported in [24] for nonlinear parameters and [25] for unobserved compartments. Beyond these method-specific issues, several general considerations apply to WENDy just as they do to forward-solver-based least squares, likelihood-based inference, Bayesian methods, gradient matching, and other approaches. In particular, WENDy does not by itself resolve model misspecification, structural or practical non-identifiability, or mismatch between the assumed and actual noise process. These concerns must still be taken into account when applying WENDy to real problems just as with any parameter estimation method.

Some more WENDy-specific limitations concern the construction of the weak-form integrals and how uncertainty quantification interacts with interpolation. The method avoids numerical differentiation by integrating against compactly supported test functions, but those integrals still need to be accurately approximated from the data. If the data is too sparse relative to the timescale of the dynamics, if fast transients occur between measurement times, or if non-uniform sampling is not interpolated carefully, the weak-form quantities can be biased. Fig. 5 illustrates this effect for sparse and noisy data: as the number of observations decreases and the noise level increases, parameter error and prediction error generally increase. Below 8 data points model parameters cannot be accurately recovered for the logistic equation, even with interpolation. Additionally, interpolated data requires particular care. While interpolation is useful for approximating quadrature when dynamics are smooth, interpolated points are not independent measurements, and treating them as independent in the WENDy covariance calculation artificially increases the effective sample size and can lead to confidence intervals that are far too narrow. For this reason, uncertainty quantification in sparse-data settings should either propagate the original measurement covariance or use a bootstrapping procedure as in Section 2.2 that resamples the original sparse data before repeating the interpolation and WENDy estimation pipeline.

We conclude by providing an outlook of the future research directions in weak form scientific machine learning (WSciML). The current frontier for WSciML methods, such as WENDy, is aimed at relaxing data and model assumptions, while preserving the main advantages of the weak-form philosophy (e.g., computational efficiency and robustness to large noise). Current research directions include a deeper exploration of nonlinear parameters in model, delay differential equations, general unobserved compartmental models, mixed-effects, switching events, and state reconstruction. Progress in these directions would make WENDy applicable to a broader class of biological models.

## 4 Methods

In this section, we present a tutorial on the Weak form Estimation of Nonlinear Dynamics (WENDy) method for parameter estimation. To have an illustrative and accessible presentation, we will introduce this method using the classical logistic equation

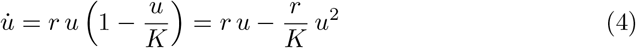

as a toy model^1^. Here, *t* ∈ [0, *T*] denotes the time variable, *u*(*t*) denotes the state variable (e.g., a population size), and *u̇* denotes its time derivative. The parameters *r >* 0 and *K >* 0 represent the intrinsic growth rate and the carrying capacity, respectively.

The WENDy framework assumes that the dynamics can be expressed as a linear combination of prescribed (possibly nonlinear) basis functions of the state. In the logistic example, we introduce *f*_1_(*u*) := *u* and *f*_2_(*u*) := *u*^2^, and rewrite (4) as

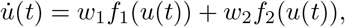

where *w*_1_ = *r* and 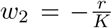. In this way, estimating the interpretable parameters (*r, K*) is equivalent to estimating the parameters (*w*_1_, *w*_2_) =: **w**^*T*^.

Suppose we observe noisy measurements of the state at uniformly spaced time points,

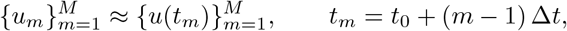

with sampling interval Δ*t >* 0.

The goal of WENDy, in this setting, is to estimate *w*_1_ and *w*_2_ (and hence *r* and *K*) from the time series 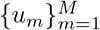 without directly approximating *u̇* from the data^2^. This is implemented by using the *weak-form* of model (4). That is, by multiplying (4) by a smooth test function *ϕ*(*t*) with compact support^3^ in (0, *T*), integrating in time, and making use of integration-by-parts, one arrives at the so called weak-form equation:

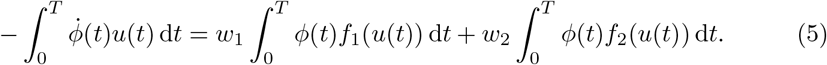

Notice that in comparison to equation (4) above, in equation (5), all derivatives have been moved from *u* to the *known* test function *ϕ*. This means that *ϕ̇* can be calculated analytically and therefore noisy approximations can be avoided. Moreover, equation is true for any smooth compactly supported *ϕ*, and so, we can use a collection of (reasonably distinct) test functions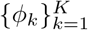 to construct a linear system of the form

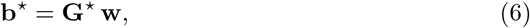

Where

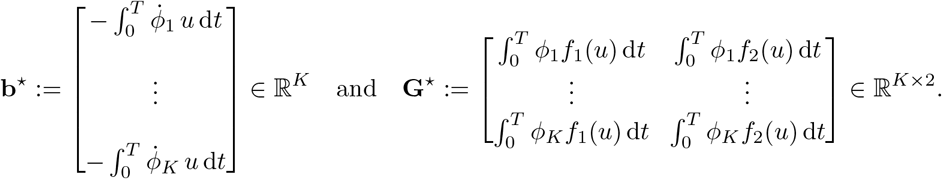

In practice, the integration in system (6) is approximated using the data, resulting in the approximated system

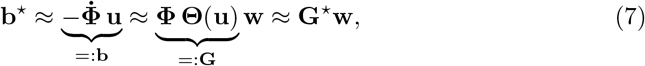

where

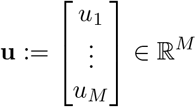

is the vector of state measurements,

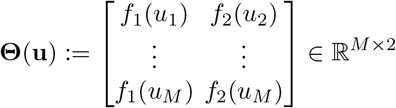

is the feature matrix evaluated at the data, and

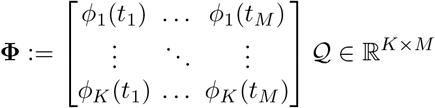

is the test function matrix evaluated at the time points multiplied by an appropriate quadrature matrix, *Q*. In our case, due to the compact support of the test function, the integration error can be mitigated by using the composite trapezoidal rule [21, Lemma 2] for which the corresponding matrix is

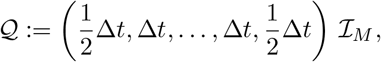

where *I*_*M*_ is the *M × M* identity matrix.

WENDy then estimates the true parameters **w** via a least squares minimization,

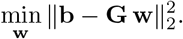

It is important to note, however, that when noise is present in the system, it appears in both the vector **b** and in the matrix **G**. Consequently, this means the least-squares problem above is an *errors-in-variables* problem and so the ordinary least-squares (OLS) estimator is generally biased, and its usual covariance formulas are invalid. The WENDy method addresses this by (i) approximating the covariance structure of the weak-form residuals and (ii) using an iteratively reweighted least squares (IRLS) procedure together with a covariance correction.

Let **r**(**u, w**) := **Gw** *−* **b** and say the observed data 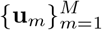 is noisy measurements of the true (noise-free) signal **u**^⋆^. The two most common noise structures are

1. **Additive normal noise:** Where 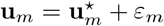 with *ε* ∼ *N* (0, *σ*^2^).
2. **Multiplicative lognormal noise:** Where 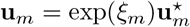.

In this case, for small noise levels, we have the first-order expansion

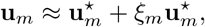

where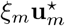 is approximately Gaussian with variance 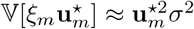.

Both cases can be handled under the assumption 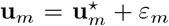 with **E**[*ε*] = 0 and Cov(*ε*) = Σ_*u*_ ∈ ℝ*M×M* which may depend on the (unknown) signal **u**^⋆^. In the additive Gaussian case, Σ_*u*_ = *σ*^2^*I*_*M*_ and is independent of the true signal **u**^⋆^. On the other hand, for the multiplicative lognormal case, the first-order approximation gives 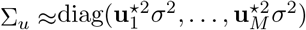, which is further approximated by substituting in the noisy measurements for the unknown true measurements, i.e., 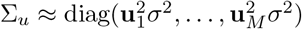 ^4^. Under this general noise formulation, the weak-form residual

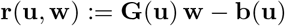

inherits noise from both **b** and **G**, since each entry represents a nonlinear functional of the noisy data **u**. A standard first-order expansion around the noise-free signal **u**^⋆^ yields

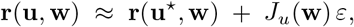

where *J*_*u*_(**w**) denotes the Jacobian of **r** with respect to the data. Consequently, the covariance of the weak-form residuals can be approximated as

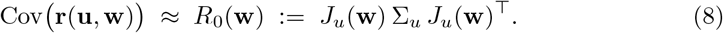

In the WENDy framework, this covariance structure is further refined by adding a second-order correction term *H*(**w**) that captures curvature effects arising from non-linear transformations of **u** (see [23] for details), so that the working covariance model is

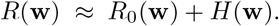

Thus, rather than solving the ordinary least-squares problem

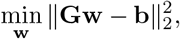

WENDy seeks to approximately solve the generalized least-squares (GLS) problem

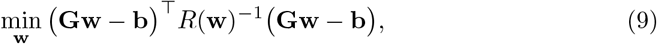

where *R*(**w**) depends on the unknown parameters through (8) and the second-order correction.

Since *R*(**w**) itself depends on **w** (and on unknown noise parameters such as *σ*^2^), WENDy employs an *iteratively reweighted least squares* (IRLS) procedure. At each iteration, the residual covariance *R*(**w**) is estimated from the current parameter iterate, factored (e.g., via a Cholesky decomposition *R*(**w**) ≈ **RR**^⊤^), and used correct autocorrelations in the residual of the regression system. The resulting transformed problem is an OLS problem:

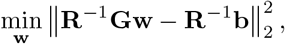

which can then be solved efficiently using standard methods. This process is repeated until convergence in **w**.

In the logistic toy problem, the WENDy algorithm proceeds by: (i) constructing the weak-form quantities **b** and **G** from the time series 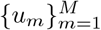, (ii) specifying a noise model (e.g., additive Gaussian or multiplicative lognormal), (iii) iteratively estimating the residual covariance *R*(**w**) and updating the parameter estimates **w** = (*w*_1_, *w*_2_)^⊤^, and finally (iv) mapping back to the interpretable parameters (*r, K*) via the identities *w*_1_ = *r* and *w*_2_ = *−r/K*.

### 4.1 Choice of test functions

Historically in the WSciML literature, there have been multiple test functions considered. For the methods presented here, we have found success two particular classes of test functions: piecewise polynomials of the form

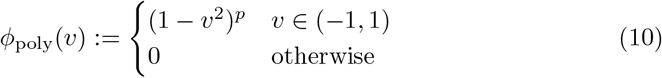

and smooth “bump” functions of the form

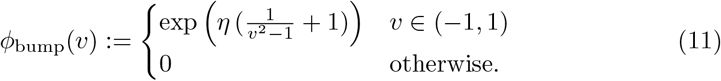

Here, *p, η >* 0 represent shape parameters which control the smoothness and decay of the test function as *v* tends to *±*1. The variable *v* := *t/r* represents a rescaling of the time variable *t* given some radius *r >* 0. This radius is a hyperparameter that is chosen from the data using the method presented in [41]. As for the shape parameters *p* and *η*, we have found that their exact value is of little consequence to the method’s performance, provided they are sufficiently large (e.g., larger than 10) to reduce the quadrature error to a negligible level. These representative test functions are then convolved with the data to construct system (7), effectively filtering out the noise while maintaining the equational relationship (5). This filtering process is demonstrated in Fig. 3(1) where we see how convolution smooths the original noisy data.

**Fig. 3.**
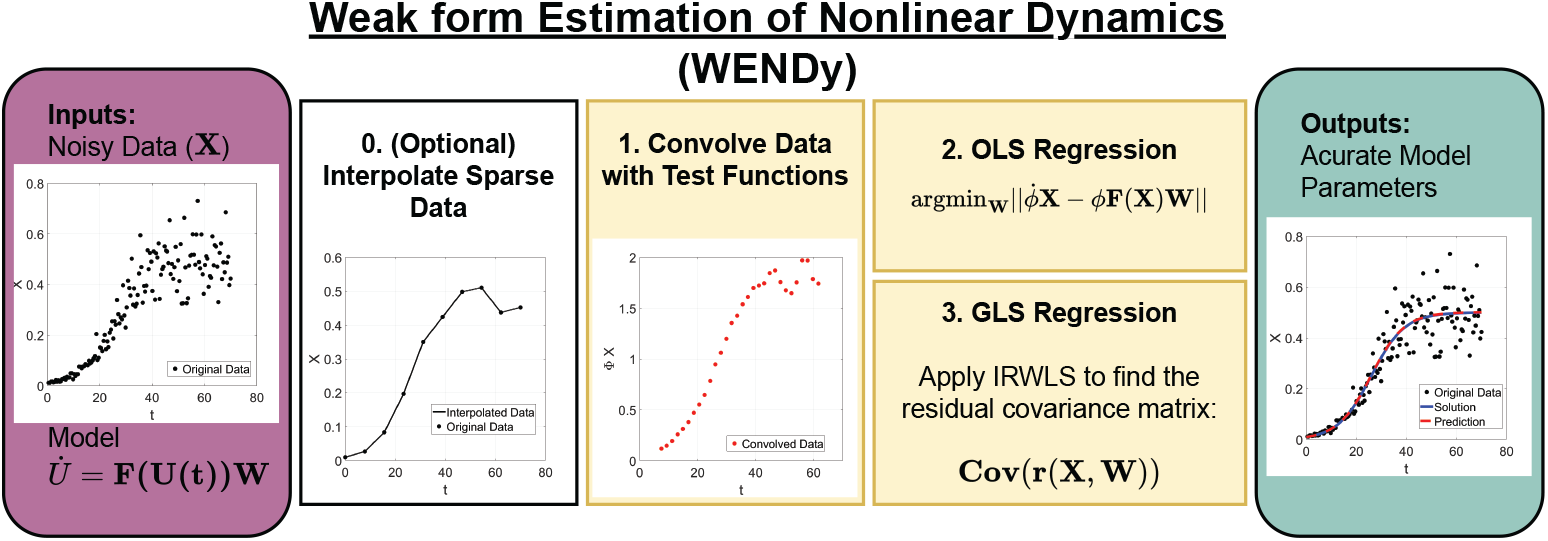
Outline of the WENDy algorithm with a simulated logistic example.

### 4.2 The case of sparse data

While traditional equation error methods are often benchmarked on chaotic, rapidly evolving systems with abundant data, many real-world applications (e.g., population growth, biochemical signaling pathways, cell-cycle progression, etc.) generate much smoother dynamics, and their experiments result in substantially sparser measurements. At first glance, this scarcity of data appears antithetical to weak-form methods, since integrating against test functions effectively reduces the number of data points as demonstrated in Fig. 3(1). However, when the underlying dynamics are smooth and slowly varying, the weak-form formulation provides significant advantages that directly counterbalance these apparent limitations.

First, the weak form entirely avoids explicit time-derivative computations. Instead, all terms in the WENDy framework appear inside integrals of the form

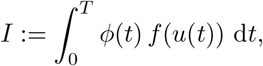

where we recall that *ϕ*(*t*) is a smooth compactly supported test function. Consequently, the data is never differentiated, and the nonlinear terms *f* (*u*(*t*)) are evaluated only at the available discrete time points where their contributions to the regression are *averaged* through numerical quadrature.

Moreover, when there are no rapid changes in the state trajectory *u*(*t*) (such as in the logistic equation), the integrand *ϕ*(*t*) *f* (*u*(*t*)) changes gradually across each test function’s support. This smoothness has an important implication: the reconstruction of *u*(*t*) between measurement times does not need to be highly accurate. As long as the interpolation respects the coarse geometry of the trajectory, the quadrature error remains small. More precisely, let *ũ* represent the piecewise linearly interpolated data onto a fine grid of spacing *h* and let

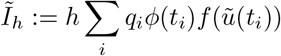

represent the composite trapezoidal rule on the interpolated data with quadrature weights 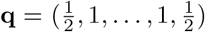. Then the total error *I − Ĩh* can be decomposed into two contributions:

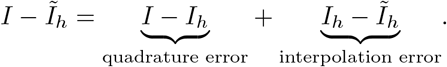

The quadrature error component inherits the fast convergence of the trapezoidal rule for periodic functions depending only on the smoothness of the integrand: approximately *O*(*h*^*p*+2^) for even *p* and *O*(*h*^*p*+1^) for odd *p* proven in [21, Lemma 2]. On the other hand, the interpolation error trends according to *O*(Δ*t*^2^) consistent with point-wise interpolation theory [42, Section 3.1]. We demonstrate these trends numerically in Fig. 4 for the noise-free logistic data and piecewise polynomial test functions from (10).

**Fig. 4.**
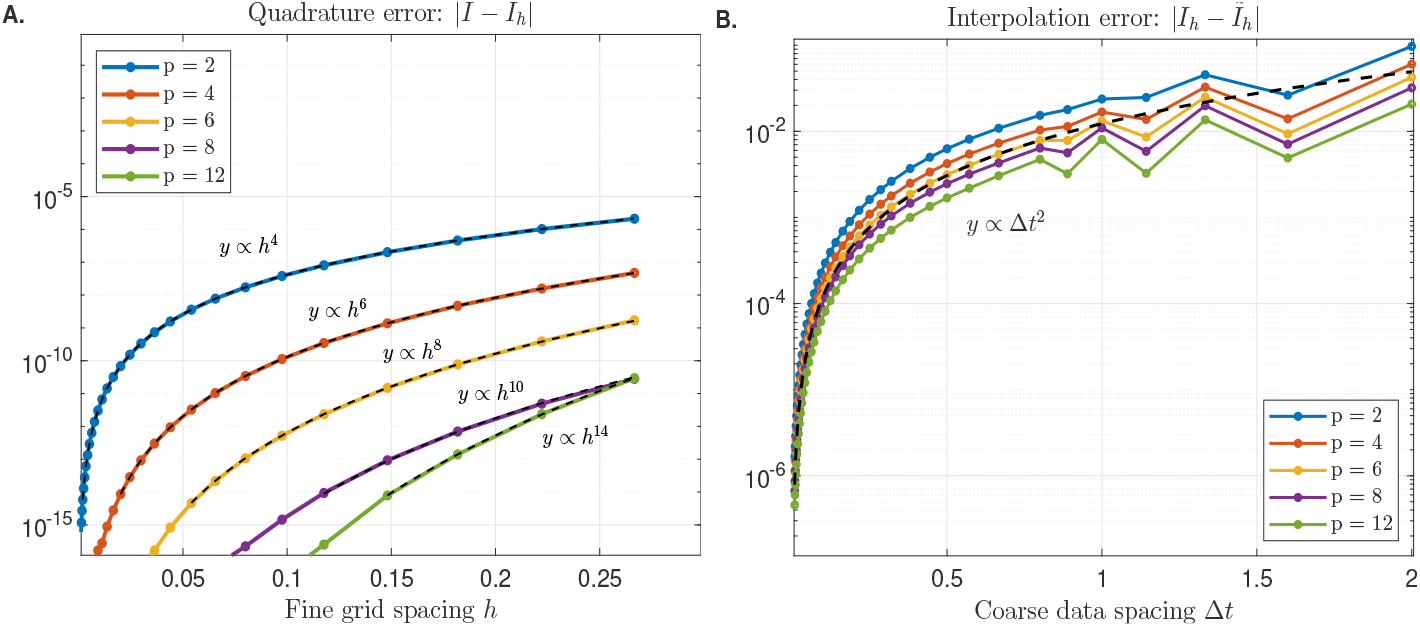
**A. Behavior of the quadrature error** | *I*− *I*_*h*_|**as a function of the interpolation spacing** *h*. **B. behavior of the interpolation error** |*I*_*h*_ − *Ĩ*_*h*_| **as a function of the sampling rate** Δ*t* **for** *h* = 10^*−*4^. The data in this plot uses the piecewise polynomial test function (10) and noise-free logistic data.

Thus, for nonchaotic systems, simple piecewise-linear interpolation of the data to “fill out” the integration quadrature is fully adequate. We test the performance of the WENDy method using linear interpolation of the data over various noise levels and number of data points in Fig. 5. Here, we study the change in the *l*^2^ parameter error metric

**Fig. 5.**
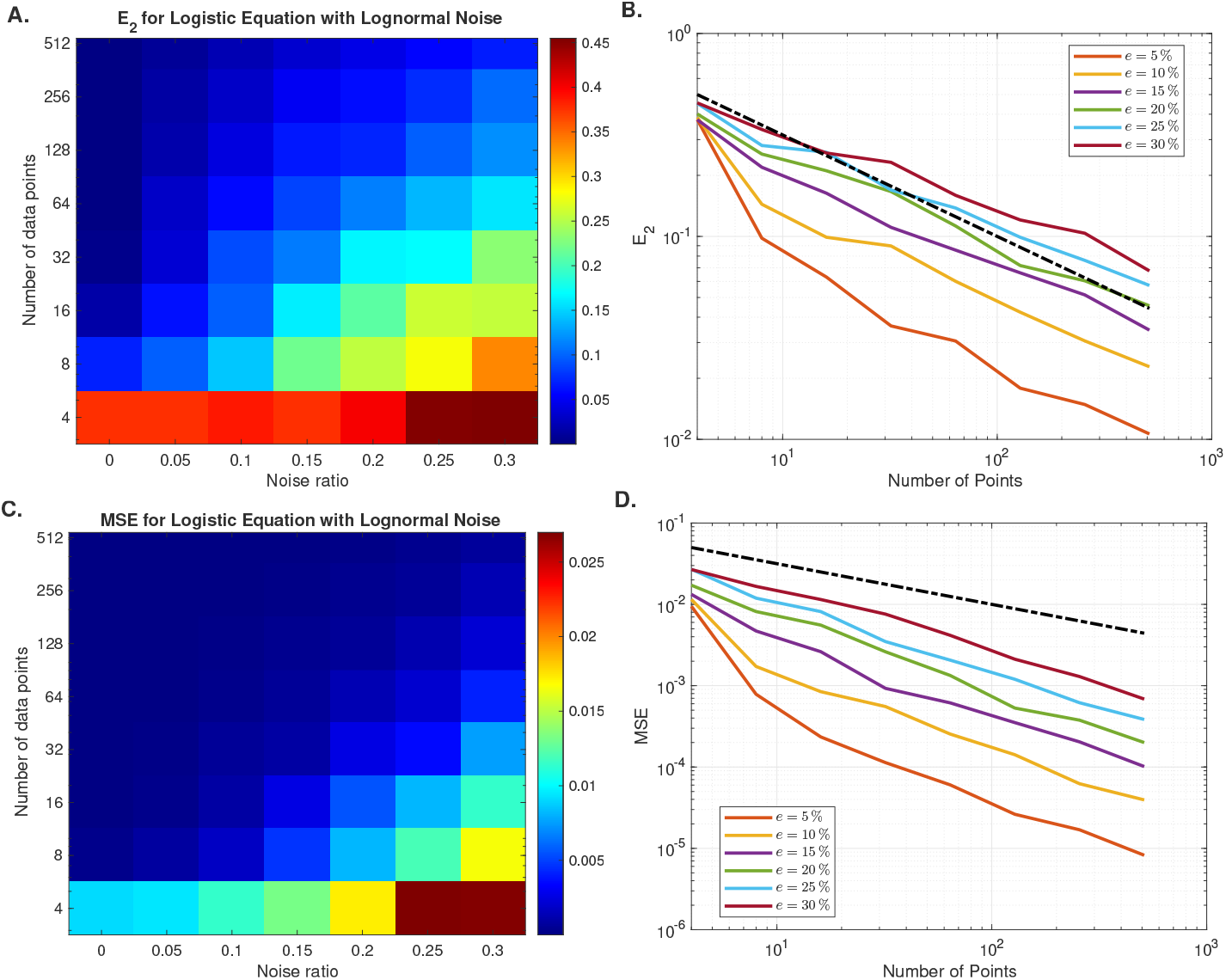
**A. Heatmap of mean** *E*_2_ **error over different numbers of data points and noise ratios. B. Trend lines of the mean** *E*_2_ **error as a function of the number of data points. Heatmap of mean mean-squared error (MSE) over different numbers of data points and noise ratios. D. Trend lines of the mean MSE as a function of the number of data points**. The black dashed line is a reference line of *O* (*N*^*−*1*/*2^). For all plots, the mean is taken over 50 realizations of the noise.

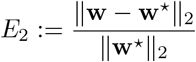

as both the number of points reduces and as the noise-to-signal *e* increases.

### 4.3 Parameter Confidence Intervals

From the residual covariance matrix, given by equation (8), we can write the approximate parameter covariance matrix^5^

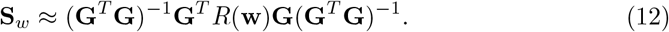

And, the (1 *− α*) confidence interval around the parameter estimate *ŵ*_*I*_

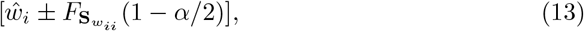

where *F* is the CDF of the normal distribution with mean zero and variance **S**_*wii*_. The estimated parameter covariance matrix is a default output of the WENDy algorithm and generally provides reliable parameter confidence intervals. Table C3 provides a comparison of the covariance approximation and parametric bootstrapping, and demonstrates that the WENDy default confidence intervals are close in magnitude to the confidence intervals generated through repeated simulation.

In the case of sparse data, we could interpret the default covariance matrix-based confidence intervals as the confidence interval if the interpolated data were the true data. In order to find the real uncertainty of the estimated parameters accounting for the interpolation step, we employ a parametric bootstrapping procedure. Briefly, we approximate the variance of the residual error using the WENDy parameter estimate. We then generate *M* = 1, 000 sets of sparse data from our assumed residual error model; recall we assume the error follows a lognormal distribution. Next, we perform the WENDy algorithm for sparse data, including a linear interpolation step, for each of the simulated data sets. Finally, we obtain the (1*− α*) confidence interval for each parameter by recovering the *α* and (1*− α*) quantiles of the set of parameter estimates. Table C4 shows a comparison of the default parameter confidence intervals versus the corrected bootstrap parameter confidence intervals for an example system with sparse data collection.

## Acknowledgments

The research reported in this publication was supported in part by the NIGMS Division of Biophysics, Biomedical Technology and Computational Biosciences (grant R35GM149335 to DMB). P.J. and M.K.J were supported by Param Hansa Philanthropies.

## Author Contributions

**N.H.B. and R.L**.: Methodology, Software, Investigation, Visualization, Validation, Writing - original draft, Writing - review & editing. **P.J**.: Data curation, Writing - review & editing. **M.K.J**.: Conceptualization, Data curation, Supervision, Writing - review & editing. **D.M.B**.: Conceptualization, Supervision, Writing - review & editing.

## Code Availability

The code used in this manuscript will be made available at the GitHub repository MathBioCU/WSciML-for-Systems-Biology.

## Appendix A Test function customization and other hyperparameters

The `wendy defaults.m` script handles the WENDy test function and solver hyperparameters, both of which could be adjusted for particular problems of interest. The defual functional form of the test function *ϕ* is the polynomial test function *ϕ*_poly_(*v*) = (1 *− v*^2^)^*p*^. We set as default *p* = 12 which results in an approximate bump function that is *C*^11^-smooth. Generally, higher values of *p* result in narrower bumps and therefore more aggressive smoothing. The script builds one `testfcn` object per trajectory and per state variable. The `‘meth’` parameter (default `‘FFT’`) tells the constructor to choose the test function’s support radius using the spectrum of the data, selecting the smallest support that captures the signal content above a threshold (see [21, 22] for a mathematical description). Each `testfcn` object precomputes and stores the convolution weights for *ϕ* and its needed derivatives, as well as the set of valid interior grid points where the test function’s support does not extend beyond the data boundaries.

Additionally, there are several hyperparameters for the WENDy algorithm whose default values are preset in `wendy_defaults.m`. The max IRLS iteration cap is set by `maxits` (defualt 100). The `ittol` parameter (default value 10^*−*4^) sets the relative convergence tolerance, i.e., the iteration terminates when

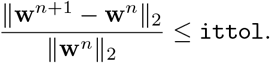

The parameter `diag_reg` (default value 0) adds a diagonal regularization to the covariance matrix before inversion, the default value being 0 disables this regularization. The boolean parameter `trim_rows` toggles outlier-trimming which removes rows of the weak-form linear system whose leverage or residual exceeds a data-driven threshold, improving robustness when individual test function placements are corrupted by boundary effects or data artifacts. Finally, `cov_params` is a 1 *×* 2 array which toggles statistical corrections in the WENDy algorithm. The first entry tells WENDy to correct for the heteroscedastic residual covariance at each iteration while the second entry tells WENDy to account for the bias introduced by noise in the feature matrix *G*.

## Appendix B Code Organization

**Table B1.** Description of the objects used in the code for this paper.

| Object | Main Responsibilities |
| --- | --- |
| `wsindy_data` | Wraps a single trajectory’s observed state vector and time grid. Stores grid metadata (spacing, uniformity, dimension) that downstream objects query when building convolution weights and determining valid index ranges. One object is created per trajectory. |
| `term` | Represents a scalar function of the state, optionally paired with a differential operator. Used to define each candidate right-hand side function and each component of the left-hand side. The optional `linOp` parameter encodes derivative order, triggering integration by parts in the weak form. |
| `addterm` | `term` subclass representing the sum of two `term` objects. Used in the creation of the left-hand side in Listing 5. |
| `library` | Collects the right-hand side `term` objects into an ordered list. Each term becomes one column of the feature matrix $\mathbf{G}$ when the model is assembled. |
| `testfcn` | Discretizes the weak-form pairing $\langle \phi, \cdot \rangle$ for a given data object. Precomputes convolution weights from the test function $\phi$ and its derivatives, determines the support radius, and identifies the valid interior indices. |
| `wendy_model` | Assembles the weak-form linear system $\mathbf{G}\mathbf{w} \approx \mathbf{b}$ from the data, library, test functions, and left-hand side. Concatenates the per-trajectory systems into a single overdetermined problem. Also stores the Hessian factors needed for iterative covariance estimation. Subclass of `wsindy_model`. |
| `WS_opt` | Solver class. Its `wendy` method runs the iteratively reweighted least squares loop: at each iteration it estimates the residual covariance $\mathbf{C}(\mathbf{w})$, whitens the system via its Cholesky factor, and re-solves the GLS problem subject to box constraints. Returns the learned weights, iterate history, asymptotic covariance, and the final Cholesky factor. |

## Appendix C Confidence Interval Generation

**Table B2.**
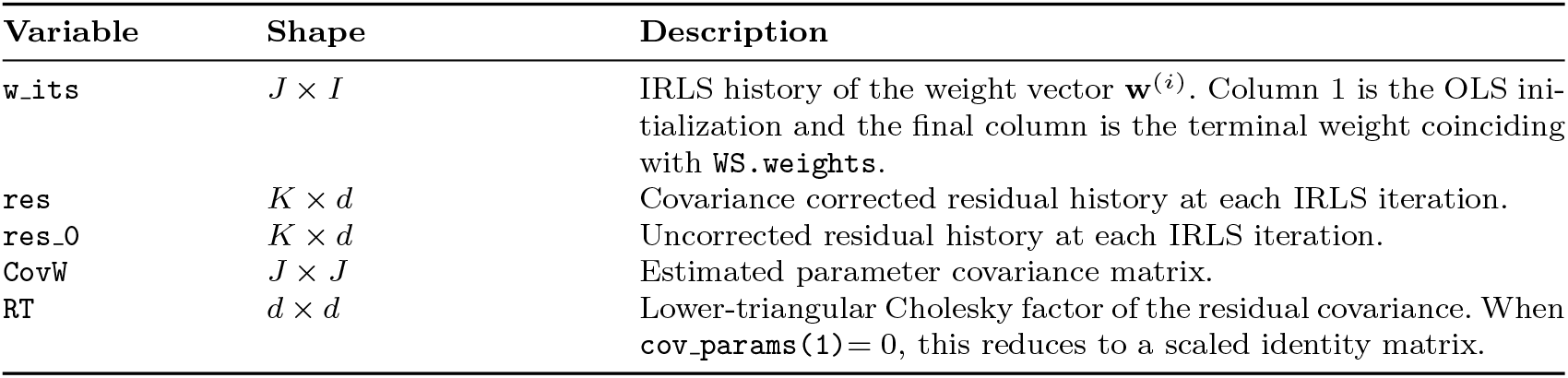
Outputs of WS Opt().wendy.

**Table C3.** Comparison of 95% confidence intervals for parameters estimated via WENDy or lsqnonlin for model (2.1) calculated with covariance matrix approximation or through parametric bootstrapping (see Section 4.3).

| Parameter Method | CI Method | $p_1$ | $p_2$ | $p_3$ | $p_4$ | $p_5$ |
| --- | --- | --- | --- | --- | --- | --- |
| WENDy | covariance matrix | [108.96, 109.04] | [35.56, 35.59] | [30.66, 30.78] | [75.36, 75.38] | [25.74, 25.77] |
|  | bootstrap | [108.95, 109.05] | [35.56, 35.30] | [30.65, 30.79] | [75.36, 75.38] | [25.74, 25.77] |
| lsqnonlin | covariance matrix | [129.72, 132.49] | [28.02, 28.60] | [64.77, 69.83] | [71.75, 72.19] | [21.85, 22.32] |
|  | bootstrap | [131.6, 137.03] | [28.18, 28.35] | [68.23, 78.65] | [71.99, 72.32] | [22.16, 22.36] |

**Table C4.** Comparison of 95% confidence intervals for parameters estimated via WENDy or lsqnonlin for models (1) and (2) calculated with covariance matrix approximation or through parametric bootstrapping (see Section 4.3).

| ParameterMethod | CI Method | CI Method | $t_m \times 10^{-2}$ | $t_e \times 10^{-2}$ | $r_e$ | $\sigma$ | $\mu$ |
| --- | --- | --- | --- | --- | --- | --- | --- |
| WENDy | G&T | covariance matrix | [7.89, 7.93] | [10.06, 10.14] | [3.3851, 3.3882] | - | - |
|  |  | bootstrap | [5.39, 15.82] | [11.08, 17.97] | [2.91, 3.59] |  |  |
|  | GI&T | covariance matrix | [6.56, 6.61] | [8.21, 8.30] | fixed | [0.496, 0.496] | [-0.2575, -0.2575] |
|  |  | bootstrap | [5.83, 15.33] | [8.28, 13.17] |  | [0.1978, 0.9689] | [-0.4076, -0.1418] |
| lsqnonlin | G&T | covariance matrix | [7.43, 7.72] | [10.03, 10.36] | [3.572, 3.5926] | - | - |
|  |  | bootstrap | [5.10, 15.08] | [3.96, 17.71] | [2.96, 4.06] |  |  |
|  | GI&T | covariance matrix | [6.38, 6.89] | [8.59, 9.23] | fixed | [0.4203, 0.4385] | [-0.2482, -0.2146] |
|  |  | bootstrap | [2.72, 13.27] | [8.76, 17.77] |  | [0.0035, 0.6850] | [-0.4265, -0.0839] |

## Footnotes

1 For a general presentation of the WENDy method for arbitrary autonomous systems of the form 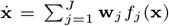, we refer the reader to the original introduction of the method [23] and the subsequent works [24, 25, 37].

2 Approximating time derivatives from noisy measurements is notoriously ill-conditioned. Finite-difference schemes amplify high-frequency noise, and even regularized or smoothing-based approaches require delicate tuning of bandwidths or penalty parameters. See, for example, classical discussions of numerical differentiation and noise amplification in [38], as well as derivative-based model discovery methods such as SINDy, which are known to degrade in accuracy under moderate noise levels unless sophisticated denoising or total-variation regularization is used [39, 40].

3 In other words, *ϕ* is only positive for values of *t* which lie in a closed sub-interval of (0, *T*).

4 In the instance of multiplicative lognormal noise and strictly positive data, one may perform the parameter identification in the log-plane using the transformation **y** = log(**u**) to avoid approximating the lognormal covariance. In this case, one would perform the parameter identification on the system 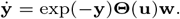

5 For a more detailed derivation of the parameter covariance matrix we direct readers to [23].

## Notes

### Competing Interest Statement

The authors have declared no competing interest.

